# Harsh snow conditions reduce body mass and reproductive success of an alpine ungulate, inducing carry-over effects

**DOI:** 10.64898/2026.05.05.723074

**Authors:** Kallan Crémel, Marco Festa-Bianchet, Alexandre Langlois, Fanie Pelletier

## Abstract

Winter can affect animal population dynamics by limiting resource availability and increasing energetic costs of movement caused by deep snow. Given the rapid alteration of snowpack properties due to climate change, quantifying how snow characteristics influence reproduction and physical condition is critical. We evaluated how snow cover duration, depth, and density affect spring body mass, reproduction probability, and subsequent autumn body mass of bighorn sheep (*Ovis canadensis*) using 45 years of individual-based data at Ram Mountain, Alberta, Canada, along with historical snow records reconstructed via the SNOWPACK model. Using Bayesian structural equation modeling, we quantified the direct and indirect effects of snow across different sex and age classes. Long and deep snow covers reduced spring body mass across all demographic groups, with yearlings, especially males, losing up to 0.12 kg per additional cm of snow depth. Harsh snow conditions reduced the probability of reproduction for adult females and generated a compensatory indirect effect on mass by avoiding the energetic costs of reproduction. In contrast, yearlings showed no compensatory responses and entered the following autumn in poor condition (up to 14% lighter for males and 8% for females following the deepest snow years). The impact of snow density on autumn mass of adult males was density-dependent, shifting from beneficial at low density (+0.09 kg per kg/m³) to detrimental at high density (-0.04 kg per kg/m³). The effects of snow conditions generate persistent, context-dependent carry-over effects across seasons. Our study suggests that distinct demographic groups rely on different mechanisms to cope with environmental constraints, highlighting complex, time-lagged consequences of changing winter climate on alpine herbivore populations.

## Introduction

Climate change has altered vegetation phenology and weather patterns (Parmesan, 2006; Parmesan & Yohe, 2003; Rosenzweig et al., 2008), particularly affecting alpine and arctic species that have limited spatial alternatives to cope with new climatic constraints (Freeman et al., 2018). In alpine ecosystems, the timing of seasons is largely driven by snow cover and temperature, which influence plant growth (Van de Kerk et al., 2018), nutrient availability and habitat accessibility (Billings & Mooney, 1968; Körner, 2003). Snow characteristics can directly affect animal survival. For instance, increased avalanche frequency or changing snow density can drastically reduce survival in mountain goats (*Oreamnos americanus*) (White et al., 2025). Climate change is expected to have both positive and negative indirect effects on animal survival and reproduction. On one hand, earlier snowmelt can extend the growing season, allowing species like yellow-bellied marmots (*Marmota flaviventris*) to increase body mass and population growth due to prolonged resource availability (Ozgul et al., 2010). On the other hand, reduced snow cover can disrupt anti-predator strategies. For example, snowshoe hares (*Lepus americanus*), whose seasonal coat color change is triggered by photoperiod, can experience a camouflage mismatch when snow melts earlier than usual: white individuals on dark, snow-free ground face increased predation, effectively turning an adaptive trait into a liability (Zimova et al., 2016).

For temperate ungulates, winter is typically the most challenging period, with limited food availability and high energetic demands for locomotion and foraging (Parker et al., 2009; Signer et al., 2011), and individuals rely heavily on somatic reserves to survive (Monteith et al., 2013). Harsh winter conditions exacerbate energy requirements, amplifying trade-offs between costly biological functions including maintenance, reproduction, and survival (Reznick et al., 2000; Williams, 1966). For instance, in reindeer (*Rangifer tarandus*), deep snow cover reduces weaning mass (Hendrichsen & Tyler, 2014), while in Alpine ibex (*Capra ibex*), shorter snow seasons result in heavier individuals (Brambilla et al., 2024). In this context, individual body mass acts as a cumulative integrator of an individual’s state, reflecting the balance between resource acquisition and energy expenditure (Monteith et al., 2014; Parker et al., 2009). Because stored energy capital is reflected in body mass, variations in mass inevitably dictate how individuals allocate resources over time, often necessitating trade-offs.

When resources are limited, trade-offs among costly biological functions can lead to carry-over effects, where events in one season influence individual performance in subsequent seasons (Harrison et al., 2011). In mammals, body mass is the primary currency mediating carry-over effects (O’Connor et al., 2014; Ricklefs & Wikelski, 2002) and act as a key determinant of fitness, influencing survival, growth, and reproduction (Ronget et al., 2018; Sauer & Slade, 1987; Stevenson & Woods, 2006). Understanding these cross-seasonal linkages is critical for predicting population resilience, particularly in stochastic environments. Because body mass integrates energy acquisition and expenditure over time, it links past environmental conditions to future performance and fitness. For instance, a long-term study on male Alpine ibex revealed that individuals with lower mass loss during winter started the growing season in better condition, leading to heavier mass the following autumn (Brambilla et al., 2024). Thus, while spring body mass is critically dependent upon winter energy expenditure, it can have carry-over influences on future mass and fitness (Pelletier et al., 2007). Pre-winter mass is crucial, particularly for adult females, as gestation occurs in winter (Parker et al., 2009; Stephenson et al., 2020). Therefore, understanding the drivers of spring body mass is essential to comprehend how environmental variation translates into demographic responses. Despite the recognized importance of winter conditions for ungulate fitness (Desforges et al., 2021; Gilbert et al., 2020), few studies have assessed the influence of snow cover characteristics on individual body mass (*e.g.,* Hansen et al.,2011) and reproduction, primarily due to the difficulty of collecting repeated measurements of body mass and fitness on known individuals.

This study aims to quantify the effects of snow conditions on spring body mass across all bighorn sheep (*Ovis canadensis*) age and sex classes. It also aims to evaluate how snow conditions interact with reproductive status in affecting the spring mass of adult females. The long-term monitoring of marked individuals at Ram Mountain, combined with historical snow records derived from the SNOWPACK thermodynamic model (Crémel et al., 2026), provides a rare opportunity to investigate the effects of snow conditions on fitness. The rut occurs between mid-November and early December (Pelletier, 2005), and gestation overlaps with the snow season. As capital breeders (Jönsson, 1997), bighorn sheep rely extensively on stored resources to meet the high energetic demands of late gestation (March-May) and early lactation (June). Consequently, mass at the onset of winter is a critical physiological baseline that determines a female’s capacity to sustain reproduction through the winter (Festa-Bianchet et al., 1998).

We used structural equation models to quantify the cascading effects of snow conditions through the life cycle, offering a causal perspective often missing from studies that rely on direct bivariate correlations. We tackled three objectives. First, we evaluated the effects of snow cover duration, depth, and density on spring body mass in both sexes, while accounting for age, autumn body mass, and population density. We expected that persistent, deep, and dense snow cover, as well as high population density, would reduce spring mass across all demographic groups. We then assessed the effects of age, autumn body mass, population density, and snow conditions on the probability of reproduction in adult females. We predicted that older females and those that were heavier in autumn would have a higher probability of reproduction, but that reproduction would be negatively influenced by high population density and harsh snow conditions. Finally, we tested for carry-over effects of snow conditions on body mass the subsequent autumn in both sexes. We expected that persistent, deep, and dense snow would reduce body mass the following autumn.

## Methodology

### Study site and population

The Ram Mountain bighorn sheep population in Alberta, Canada, has been the subject of a comprehensive long-term wildlife monitoring effort since 1971 (Festa-Bianchet et al., 2017). The study site is approximately 30 kilometers east of the main Canadian Rocky Mountains (52°20’50.2 N; 115°46’09.2 W). It includes 38 km² of subalpine meadows and alpine tundra (Jorgenson et al., 1993). Elevations used by sheep range from 1430 to 2160 m above sea level (Cloutier et al., 2024). The population is demographically isolated with minimal immigration or emigration (Festa-Bianchet et al., 2019), allowing for precise demographic tracking.

### Sheep capture and data collection

We captured sheep annually between late May and late September in a corral trap baited with salt. At each capture, we measured body mass with a Detecto spring scale (± 250 g). On average, individuals were weighed 3.53 times per season (SD = 1.42) (Larue et al., 2022). Body mass was standardized to June 5 and September 15 each year using mixed-effects models (Martin & Pelletier, 2011). Repeated measurements of body mass of the same individual within and between years allow the assessment of seasonal and interannual changes in body mass from birth to death.

Female reproductive status was evaluated at capture based on the presence of milk or colostrum. Because bighorn ewes can conceive as yearlings (Festa-Bianchet et al., 2019; Jorgenson et al., 1993), our analysis included all females aged one year or older in autumn (year *t*). Sheep were therefore two years or older when their spring body mass was recorded in year *t+1*. Population density was defined as the number of ewes aged ≥ 2 years present in June (Cloutier et al., 2024) (range 16 - 103). We analyzed 45 years (1979-2024) of data for adults, and 44 years (1980-2024) for lambs that survived their first winter and were yearlings at the time of spring weighing. All animal procedures were approved by the Université de Sherbrooke Animal Care Committee (Protocol FP: 2024-4569) in accordance with the guidelines of the Canadian Council on Animal Care.

### Snow characteristics

We used a historical reconstruction of snow characteristics since 1979. Complete methodology and validation for this dataset are available in Crémel et al. (2026). Briefly, historical snow data were reconstructed using simulations from SNOWPACK (Bartelt & Lehning, 2002; Lehning, Bartelt, Brown, & Fierz, 2002; Lehning, Bartelt, Brown, Fierz, et al., 2002), a thermodynamic snow model, and forced using calibrated meteorological data from the North American Regional Reanalysis (Mesinger et al., 2006). This reconstruction provided several metrics each snow season: duration of snow cover (days); mean, median and maximum depth (cm); density (kg/m^3^) and snow water equivalent (mm); and the start and end dates of snow cover each year. As several of these metrics were highly correlated (*r* > 0.6), for subsequent analyses, we only retained a subset that was weakly or not correlated and directly linked to our hypotheses: duration of snow cover, mean snow depth and median density (Appendix S1: Figure S1).

### Statistical analysis

We used Bayesian Structural Equation Models to assess causal pathways (Yuan & MacKinnon, 2009) between snow characteristics, population density, reproductive status, and individual mass. We performed all analyses using *R 4.5.1* (R Core Team, 2025). Prior to modeling, we standardized (z-scored) continuous explanatory variables (population density, snow depth, duration, density, and sheep age) to facilitate effect size comparison and algorithm convergence. For adults, age was modeled using natural cubic splines in the *splines* package (Bates & Venables, 2022) with three degrees of freedom (*df* = 3) to account for non-linear patterns in mass and reproduction probability (Appendix S1: Table S2).

To disentangle the direct and indirect effects of snow characteristics and population density on body mass dynamics and reproduction, we employed a multivariate Bayesian multilevel modeling framework. This approach allowed us to estimate parameters for multiple response variables simultaneously while propagating uncertainty across the entire causal pathway.

We constructed three distinct model sets based on demographic group. This separation was motivated by the marked sexual differences in age-specific mass, and by the distinct mass trajectories of lambs compared to older age classes (Festa-Bianchet et al., 1996). Modeling these groups separately avoided overly complex interaction terms and simplified the causal structure of the models (Figure 1):

1. Lambs: A two-equation model predicting (i) spring body mass (*t+1*) and (ii) body mass the subsequent autumn as yearlings (*t+1*).
2. Females: A three-equation model predicting (i) probability of reproduction (*t+1*), (ii) spring body mass (*t+1*), and (iii) subsequent autumn body mass (*t+1*).
3. Males: A two-equation model predicting (i) spring body mass (*t+1*) and (ii) subsequent autumn body mass (*t+1*).

**Figure 1:**
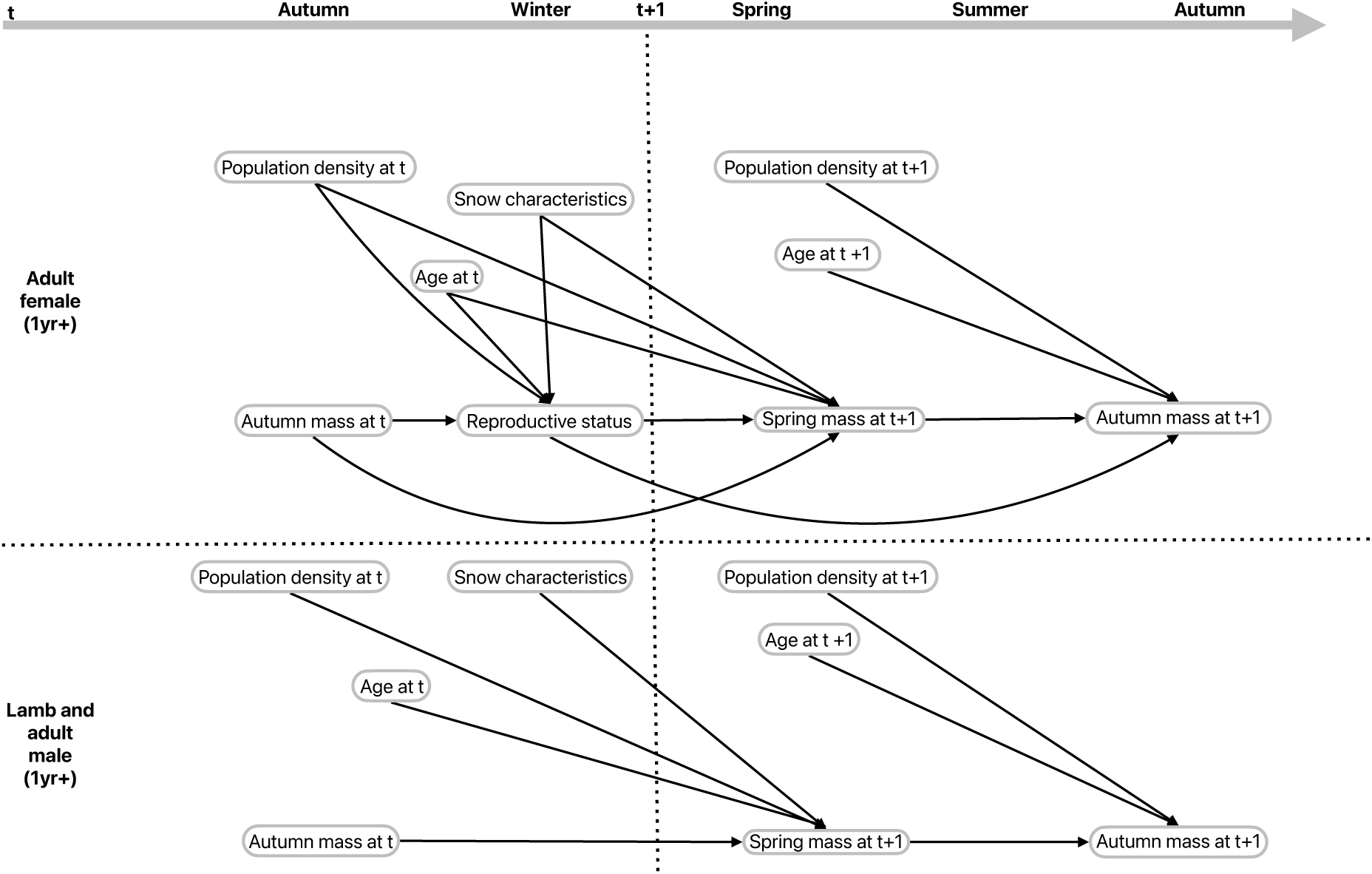
Conceptual path diagram of the structural equation model (SEM) assessing the effects of snow characteristics, age, and population density on seasonal body mass dynamics of bighorn sheep on Ram Mountain, Alberta, Canada. The model is stratified into two groups: adult females (top panel) and yearling/adult males (bottom panel). Arrows represent hypothesized causal relationships spanning from autumn *t* to autumn *t+1*. Snow characteristics and population density are modeled to affect spring body mass directly. Autumn body mass at *t* is modeled as a determinant of spring body mass at *t+1*, which subsequently influences autumn body mass at *t+1*. For adult females, reproductive status at *t* acts as a mediating variable, linking autumn body mass and age to spring body mass.

In the model for lambs, we included an interaction between sex and mass, as well as sex and environmental covariates, to account for the sexual difference in growth rates and higher sensitivity of males to the environment compared to females (Festa-Bianchet et al., 1996; Leblanc et al., 2001). For adult females, we modeled an interaction between reproductive status and spring mass. As gestation and lactation are energetically costly, reproductive females typically gain less mass and enter autumn in poorer body condition compared to non-reproductive ones (Festa-Bianchet, 1998).

We fitted all models using the *brms* package (Bürkner, 2017) version 2.22. We used set_rescor = FALSE to assume independent residuals across equations. We used default weakly informative priors for all parameters. We ran the models with four chains and 6,000 iterations per chain (including 2,000 warm-up), confirming convergence both visually and through R-hat values (all = 1.00). Effective sample sizes for both bulk and tail exceeded 400, following recommendations from Vehtari et al. (2021). We assessed model fit via posterior predictive checks (see Appendix S2: Figure S1 to S7). To investigate the effects of snow characteristics on spring body mass and the effect of spring mass on mass the following autumn, models included year and individual identity (ID) as random intercept to account for repeated measurements and inter-annual variation. For models restricted to lambs, we included only year as a random intercept. We used models with a Bernoulli distribution, including random intercept for year and ID, to assess effects on the probability of reproduction. We defined ‘year’ in the model as the calendar year in which each predictor variable was measured.

For each demographic group, we compared a candidate set of models (Appendix S1: Table S1 & S3). Model performance was assessed using K-fold Cross-Validation (K=10). We compared models based on the Expected Log Predictive Density (ELPD) and retained the model with the highest predictive performance. To ensure that these default priors did not unduly influence posterior estimates, we performed a power-scaling sensitivity analysis using the *priorsense* package, version 1.2 (Kallioinen et al., 2023). We assessed the sensitivity of the posterior distributions to perturbations in both the prior and likelihood based on the cumulative Jensen-Shannon distance. This analysis confirmed that our results were primarily driven by the data rather than the prior distributions (Appendix S1: Table S4).

When models contain interactions, the interpretation of main effects alone can be misleading as they often mask context-dependent responses. To dissect these interactions, we applied two complementary approaches based on the nature of the variables involved. For interactions including a categorical variable (*e.g.*, *Reproduction status × Average snow depth*), we performed a conditional marginal trend analysis (Lenth, 2016) to quantify and compare the slope of the environmental effect within specific biological contexts (*e.g.*, reproductive vs. non-reproductive females). For interactions involving two continuous variables, we employed the Johnson-Neyman technique (Johnson & Neyman, 1936). Unlike conventional marginal trend analyses that test significance at arbitrary points (*e.g.*, mean ± 1 SD), the Johnson-Neyman technique can determine the regions of significance and identify the range of values of the moderator variable within which the slope of the environmental driver is different from zero.

To quantify how snow conditions affect body mass the subsequent autumn (*t+1*), we performed posterior predictive simulations based on the selected models. The choice of the causal inference method was driven by the structural complexity of each model. For models without mediating bifurcations, we quantified the cumulative effect of snow conditions. We simulated the propagation of an environmental change (*e.g.*, +1 cm of snow depth) through the causal chain. The predicted difference between the “high snow” simulation and a reference simulation (mean individual in a mean environment) represents the net carry-over effect.

For models involving mediating bifurcations (*e.g.*, with Reproduction status), we employed a counterfactual mediation framework (Imai et al., 2010; Pearl, 2021). This approach relies on simulating specific counterfactual simulations, where individuals experience the reproductive conditions of a “high snow” year but theoretically retain the body mass they would have achieved during a mean snow year. This allows us to decompose the total effect of snow conditions into two distinct pathways. First, we estimated the indirect effect via body mass, representing snow effects on mass accumulation, independent of reproduction. Second, we estimated the indirect effect via reproduction, representing the mass change caused solely by the shift in reproductive success. The sum of these two distinct pathways constitutes the total effect. A comprehensive visualization of this procedure is provided in Appendix S1: Figure S2.

We also reported the probability of direction (pd) to assess the certainty of trends. The pd is a Bayesian metric defined as the proportion of the posterior distribution that shares the same sign as the median (Makowski, Ben-Shachar, & Lüdecke, 2019; Makowski, Ben-Shachar, Chen, et al., 2019). A pd of 95% indicates a 95% probability that the effect is positive (or negative), providing a direct and intuitive measure of effect presence. To visualize the magnitude of indirect effects, we employed the prediction-based framework proposed by Bush-Beaupre et al. (2026). Unlike traditional metrics that solely report a coefficient, this method illustrates the indirect pathway by superimposing the predicted shift in the mediator variable (driven by environmental exposure) onto the response curve of the outcome variable. The resulting highlighted segment on the curve isolates the specific change in the outcome attributable solely to the environmentally induced variation in the mediator, allowing for an intuitive decomposition of complex causal chains.

## Results

### Spring body mass

Body mass the previous autumn was the primary driver of spring body mass across all demographic groups (Appendix S3: Table S1). The best-supported model for yearlings was the base model (Appendix S1: Table S1 and S5). Average snow depth negatively influenced yearling spring mass (Appendix S3: Table S1, Figure 2A), with a decrease of 0.08 ± 0.03 kg (SE) for each additional centimeter of snow for females. While the 95% credible intervals slightly overlap 0 for the interaction between snow depth and sex, the data suggested a stronger negative response to average snow depth for males (pd = 95%) than for females with a decrease of 0.12 ± 0.03 kg (SE) for each additional cm of snow (Figure 2A). Snow cover duration reduced spring mass by 0.03 kg for each additional day of snow cover (Appendix S3: Table S1, Figure 2B). Effects of the other snow variables on sex-specific body mass remained uncertain overall (pd of 63% for snow median density and 53% for the interaction between sex and snow duration). In contrast, we found an interaction between sex and population density, indicating a negative effect of population density on spring mass for males but not for females (Appendix S3: Table S1). Aside from this interaction, the data did not provide strong evidence for a directional main effect of population density (pd = 57%) or median snow density (pd = 53%) on yearling spring mass, as their 95% credible intervals broadly overlap zero (Appendix S3: Table S1).

**Figure 2:**
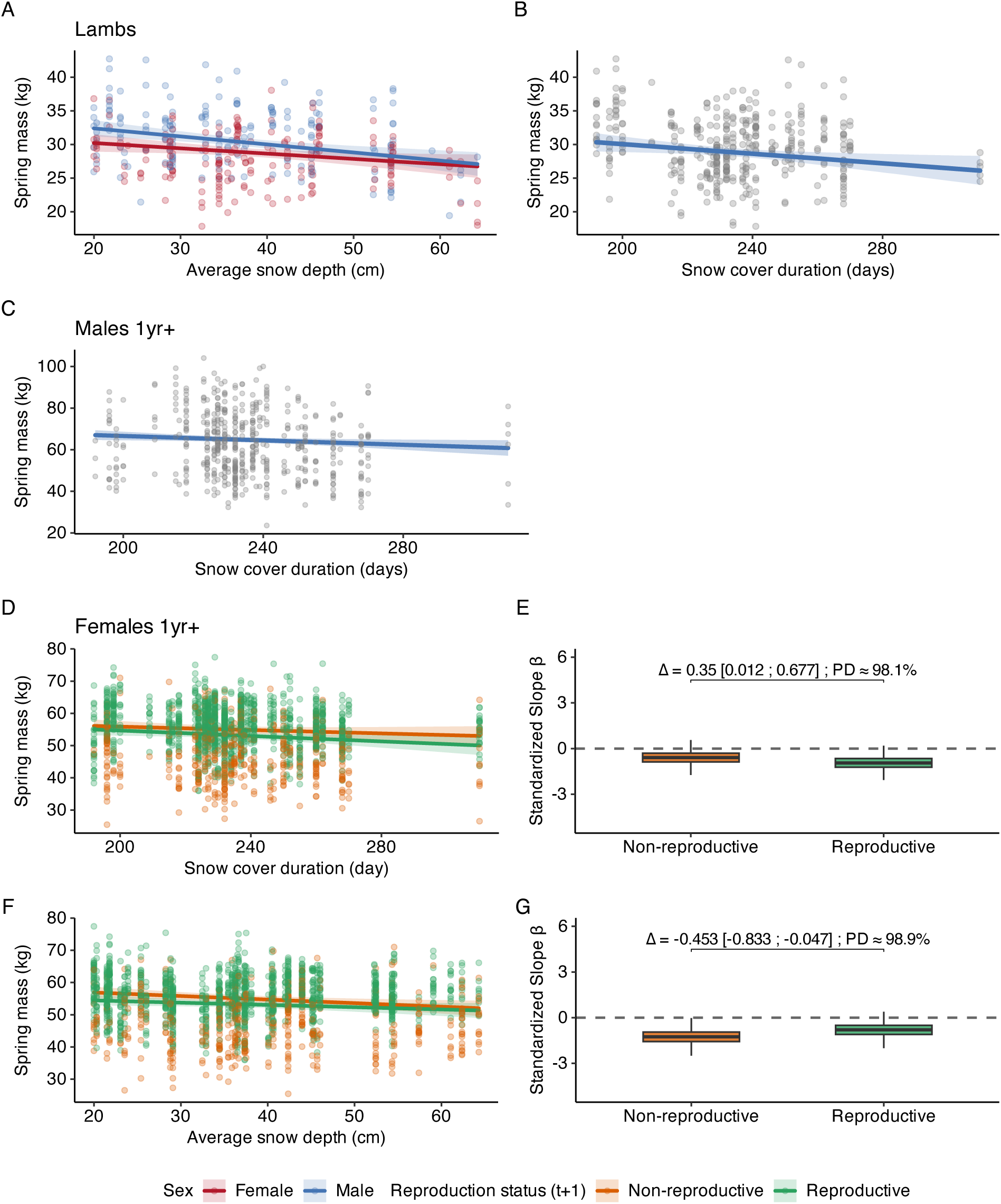
Panels A–C show the effects of average snow depth (cm) displayed by sex (red: female, blue: male) and snow cover duration (days) on the spring mass of lambs that survived their first winter (A, B; n = 373) and of males aged 1 year or older in year *t* (C, n = 514 over 45 years). The lines and shading represent fitted linear predictions with 95% credible intervals (CI). Panels D–G display results for females aged 1 year or older in year *t* (n = 1,746 over 45 years). D and F show raw data points colored by reproductive status at *t+1* (orange: non-reproductive; green: reproductive), with fitted predictions for snow cover duration (D) and average snow depth (F). Panels E and G present the posterior distributions of standardized slopes (ß) describing the effects of snow depth and cover. Boxes represent the interquartile range, horizontal lines indicate medians, and whiskers show the 95% CI. Dashed horizontal lines indicate no effect (ß = 0). Values above brackets (Δ) correspond to the difference in slopes between reproduction status, with associated 95% CI and probability of direction (PD).

For adult females, the best model included reproductive status (Appendix S1: Table S1 and S5). Females that had given birth were ∼1.8 kg lighter than non-reproductive females the following spring (Appendix S3: Table S1). We found no strong directional effect of median snow density on adult female spring mass (pd = 79%), and no evidence of an interaction with reproductive status (pd = 50%) as both credible intervals substantially overlapped 0 (Appendix S3: Table S1). All females lost mass as snow cover duration increased, but reproductive females consistently exhibited lower spring body mass across all durations (Appendix S3: Table S1, Figure 2D). Comparing the slopes confirmed that snow cover duration affected reproductive females more severely (β _repro_ = -0.947, 95% CI [-1.76; -0.11], pd = 99%) than non-reproductive females, for whom the trend was weaker and more uncertain (β _non-repro_ = -0.592, 95% CI [-1.43; 0.27], pd = 91%; Figure 2E). We also found an interaction between reproductive status and average snow depth (Appendix S3: Table S1, Figure 2F). Increasing snow depth reduced spring mass for both groups, but the effect was more pronounced for non-reproductive females (β _non-repro_ = -1.258, 95% CI [-2.19; -0.36], pd = 99%) than for reproductive females (β _repro_ = -0.802, 95% CI [-1.69; 0.07], pd = 97%; Figure 2G). Overall, non-reproductive females weighed on average ∼1.6 kg more than reproductive females in autumn.

The best-supported model for adult males included an interaction between median snow density and population density (Appendix S1: Table S1 and S5). Consistent with results from other groups, autumn body mass positively affected spring mass (Appendix S3: Table S1), while snow duration (Appendix S3: Table S1, Figure 2C) exerted negative effects on mass the following spring. Although models also evaluated interactions between population density and other snow metrics, the interaction of snow duration was highly uncertain (pd = 67%), whereas the interaction with snow depth was more certain (pd = 87%). The interaction between median snow density and population density (Appendix S3: Table S1, Figure 3A) revealed a density-dependent response (pd = 98%): the effect of snow density on spring mass was positive (Figure 3B), and the credible interval was different from zero at densities of ∼15 to 35 individuals. However, as population density increased past ∼40 females, the 95% credible interval included zero. At the highest observed densities, the slope trended to negative, suggesting that the beneficial effect of dense snow may reverse at high density. Similarly, the interaction between snow depth and population density (Appendix S3: Table S1, Figure 3C) revealed a density-dependent response with a negative effect of snow depth on spring mass that weakened as density increased (Figure 3D), with the 95% credible interval crossing zero at approximately 76 individuals. This suggests that the detrimental effect of deep snow may be reduced at the highest population density.

**Figure 3:**
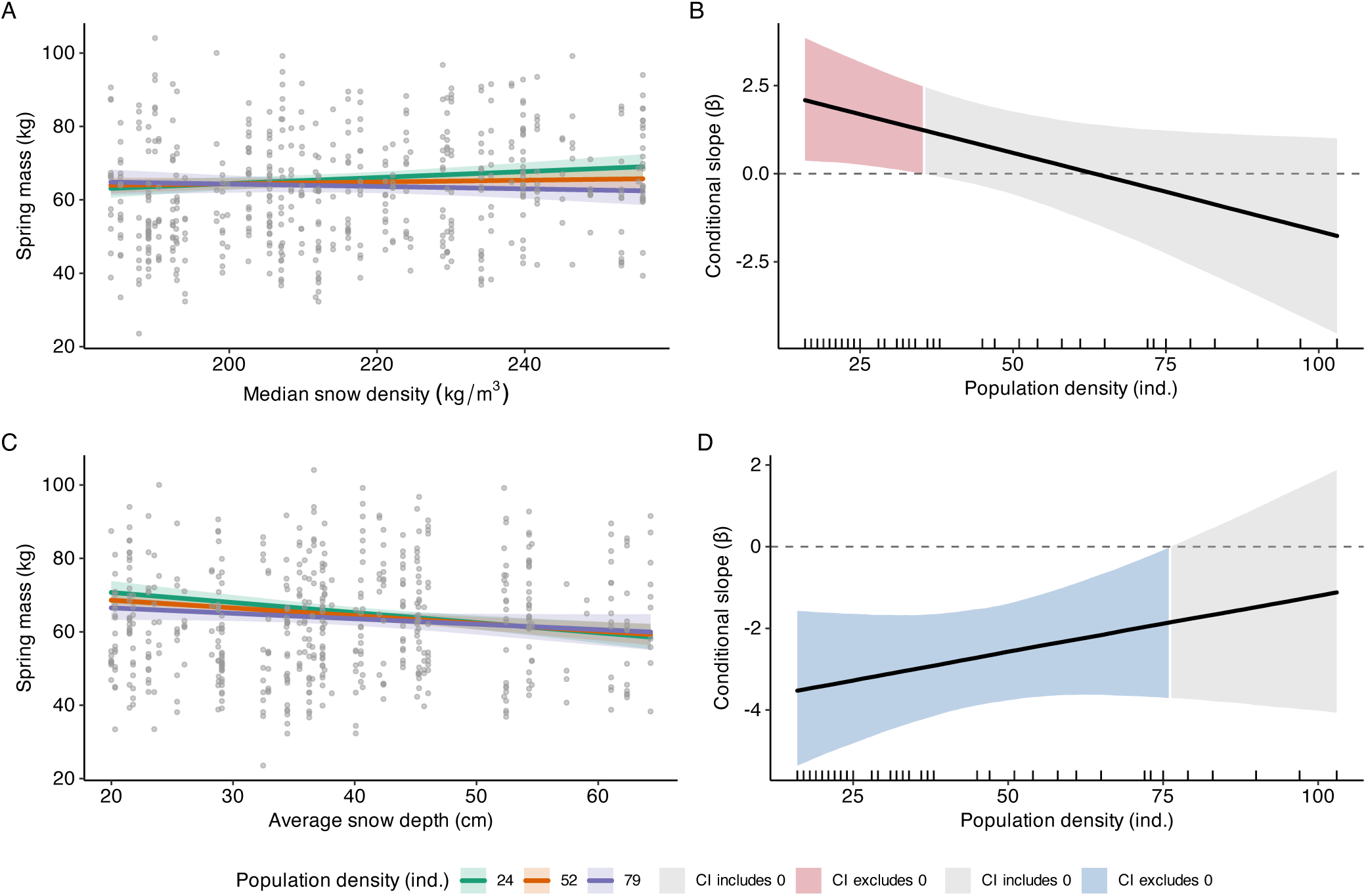
(A) Relationship between spring body mass (kg) of adult bighorn sheep males on Ram Mountain, Alberta, Canada, and median snow density (kg/m³) at three levels of population density (low = 24 individuals, green; mean = 52, orange; high = 79, purple). Points represent raw data, with fitted linear predictions and 95% credible intervals. (B) Johnson-Neyman plot illustrating the conditional slope (ß) of snow density on spring body mass across the continuous gradient of population density. (C) Relationship between spring body mass (kg) of adult bighorn sheep males and average snow depth (cm) at three levels of population density. Points represent raw data, with fitted linear predictions and 95% credible intervals. (D) Johnson-Neyman plot illustrating the conditional slope (ß) of average snow depth on spring body mass across the continuous gradient of population density. The solid black line represents the estimated change in slope, and the shaded area indicates the 95% credible interval. The red or blue areas highlight the range of population density where the credible interval of snow density or snow depth excludes 0, while the grey area indicates where the credible interval includes 0. Vertical ticks along the x-axis represent the distribution of population density observations. Based on 514 observations over 45 years.

### Offspring production

The likelihood to produce a lamb the following spring increased with female age (Appendix S3: Table S1), with a sharp increase between ages 1 and 3, peaking at 90-95% probability of reproduction between ages 3 and 10, followed by a gradual decline thereafter (Figure 4C). Reproductive probability was also strongly positively influenced by body mass the previous autumn (Appendix S3: Table S1), with at least a 95% chance of reproducing from 64 kg upwards (Figure 4D). Both snow cover duration and average snow depth had negative effects on reproduction probability, with longer lasting and deeper snow cover associated with lower reproduction (Appendix S3: Table S1 & Figure 4A & B). There was no evidence of a directional effect of median snow density on adult female reproductive success (pd = 53%) (Appendix S3: Table S1).

**Figure 4:**
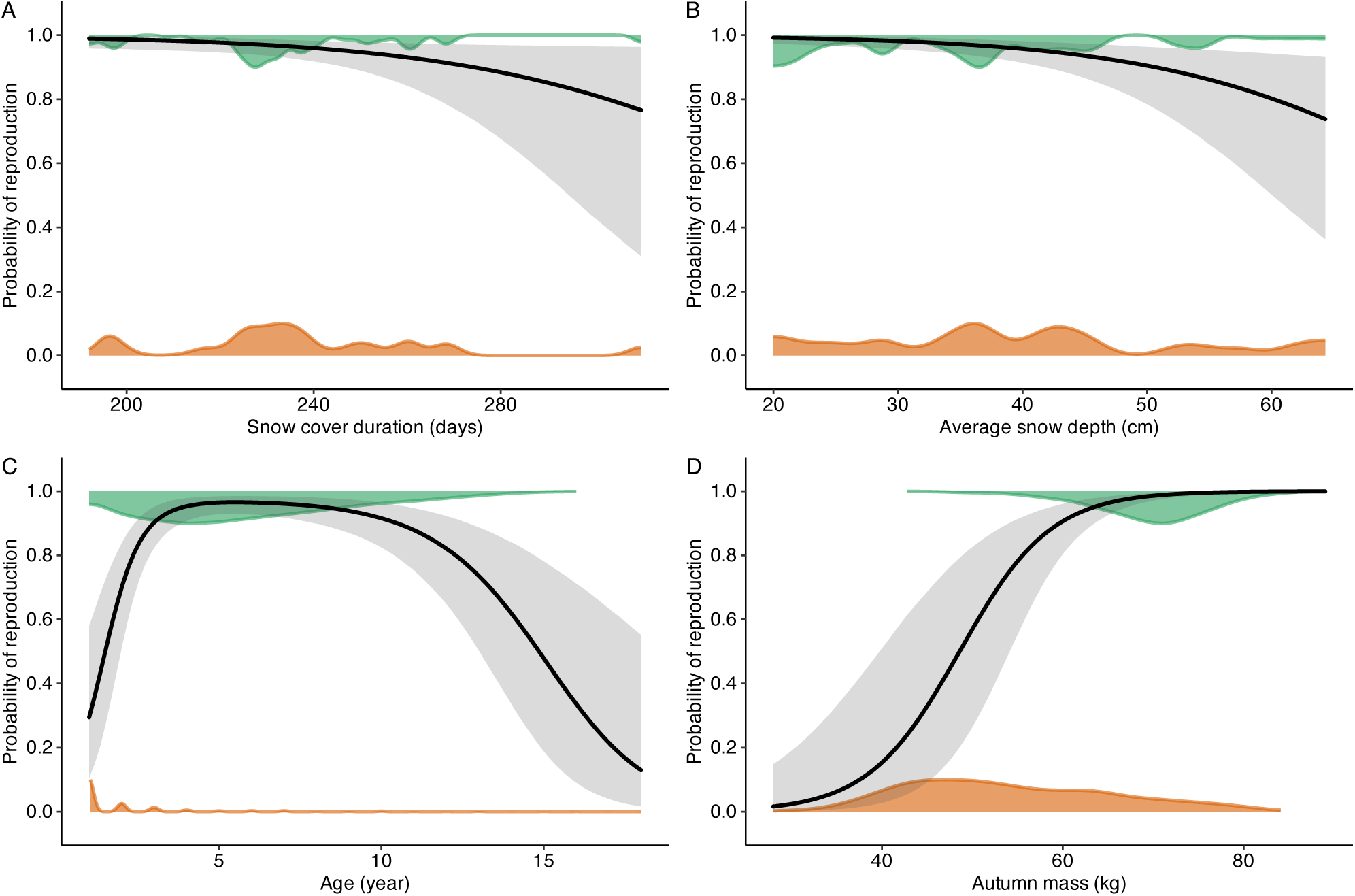
Effects of snow cover duration (A), depth (B), age (C) and autumn body mass before conception (D) on reproductive probability in year *t+1* for female bighorn sheep aged 1 year and older. The distributions (orange: non-reproductive and green: reproductive) represent the raw data plotted against each explanatory variable, with fitted linear predictions (black lines) and 95% credible intervals (dark gray). Based on 1746 observations over 45 years.

### Autumn mass and carry-over effect

Regardless of snow conditions, spring body mass strongly predicted body mass the following autumn across all demographic groups (Appendix S3: Table S1). Heavier individuals at the end of winter remained the heaviest in the subsequent autumn. Reproductive females were lighter than non-reproductive ones (Appendix S3: Table S1) in autumn. Male yearlings tended to be heavier than female yearlings (Appendix S3: Table S1).

For lambs that survived the winter, snow conditions (*t*) had a carry-over effect on their autumn body mass (*t+1*) as yearlings. An increase of 1 cm in snow depth resulted in a net loss of 0.08 kg (95% CI: -0.15; -0.02) for females and 0.15 kg (95% CI: -0.23; -0.08) for males (Figure 5A). Across the observed range of snow depth, this translated into a substantial difference: ∼6.7 kg for males (-13.7% of mean mass) and ∼3.7 kg for females (-7.9% of mean mass) following the years with the deepest and shallowest snow (Figure 5B). Similarly, an increase of 1 day of snow cover reduce mass by approximately 0.04 kg for females (95% CI: -0.07; -0.01) and 0.05 kg for males (95% CI: -0.08; -0.01) (Figure 5C). Comparing the shortest and longest snow cover observed reveals a loss of ∼5.5 kg for males (-11% of mean mass) and ∼4.5 kg for females (-9.7% of mean mass) (Figure 5D). In contrast, median snow density did not affect yearling autumn body mass, with an increase of 1 kg/m³ resulting in an extremely uncertain effect for both females (-0.001 kg, 95% CI [-0.04; 0.04], pd = 52%) and males (0.004 kg, 95% CI [-0.04; 0.05], pd = 59%).

**Figure 5:**
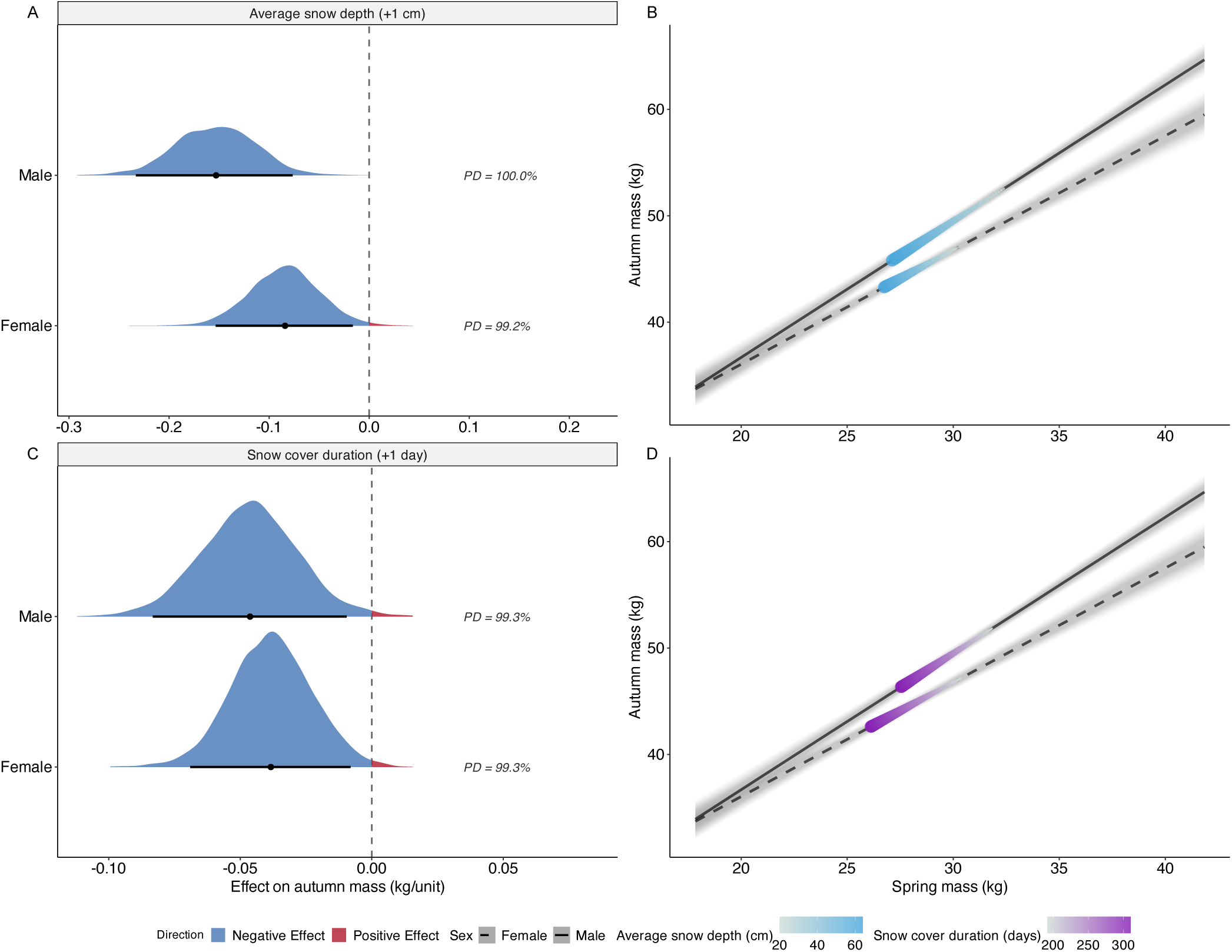
Carry-over effects of snow conditions on subsequent autumn body mass in bighorn sheep yearlings. Left panels show the posterior distributions of the indirect effect of (A) average snow depth (cm) and (C) snow cover duration (days) on autumn body mass. Estimates are show by sex (males: top; females: bottom). Density ridges represent the full posterior distributions, colored by direction (blue indicates a negative effect and red a positive effect). Black points indicate the posterior median, and horizontal bars show 95% credible intervals. The vertical dashed line denotes no effect (0), and values indicate the probability of direction (PD). Right panels show the visualization of the carry-over effect magnitudes following Bush-Beaupre et al. (2026). Regression lines show the effect of spring body mass on autumn body mass (males: solid line; females: dashed line). The colored segments (blue: snow depth; purple: snow cover duration) illustrate the predicted indirect pathway of snow conditions on autumn mass. These ribbons isolate the specific change in mass strictly induced by snow variables. Reading the segments along their color gradient illustrates the indirect effect: as snow conditions become more severe, the impact induced on spring mass (the shift along the x-axis) subsequently dictates a proportional change in autumn mass (shifting down on the y-axis). The convex (bulging) side of the colored ribbon indicates the directionality of the effect.

In adult females, the effects of previous snow conditions on autumn body mass were complex. While the total effect of average snow depth and duration was negative (ß _Total_ = -0.03 kg/cm 95% CI [-0.07; 0.01] and -0.02 kg/day 95% CI [-0.04; 0.001] (Figure 6A & C)), these trends were a bit more variable compared to those found for yearlings (Snow depth: pd = 93%; Duration: pd = 97%). This partly arises because the negative effect mediated through spring mass was partially offset by a positive indirect effect via reproduction: females experiencing harsher snow conditions were less likely to reproduce, thereby gaining more mass by autumn. The net reduction in autumn mass between the deepest and shallowest snow was around 1.63 kg (-2.4% of mean mass) for reproductive females and around 3.1 kg (-4.4% of mean mass) for non-reproductive ones (Figure 6B). Likewise, the net reduction in autumn mass between the years with the longest and shortest snow cover duration was 2.62 kg (-3.9% of mean mass) for reproductive females and 1.97 kg (-2.8% of mean mass) for non-reproductive ones (Figure 6D). Finally, median snow density had no effect on adult female autumn mass. An increase of 1 kg/m³ yielded a negligible and highly uncertain indirect total effect (ß _Total_ = 0.01 kg, 95% CI [-0.01; 0.03], pd = 79%), with similarly non-significant indirect effects mediated through spring mass (ß _Mass_ = 0.01 kg, 95% CI [-0.01; 0.03], pd = 79%) and reproduction (ß _Repro_ <0.01 kg, 95% CI [-0.00; 0.00], pd = 53%).

**Figure 6:**
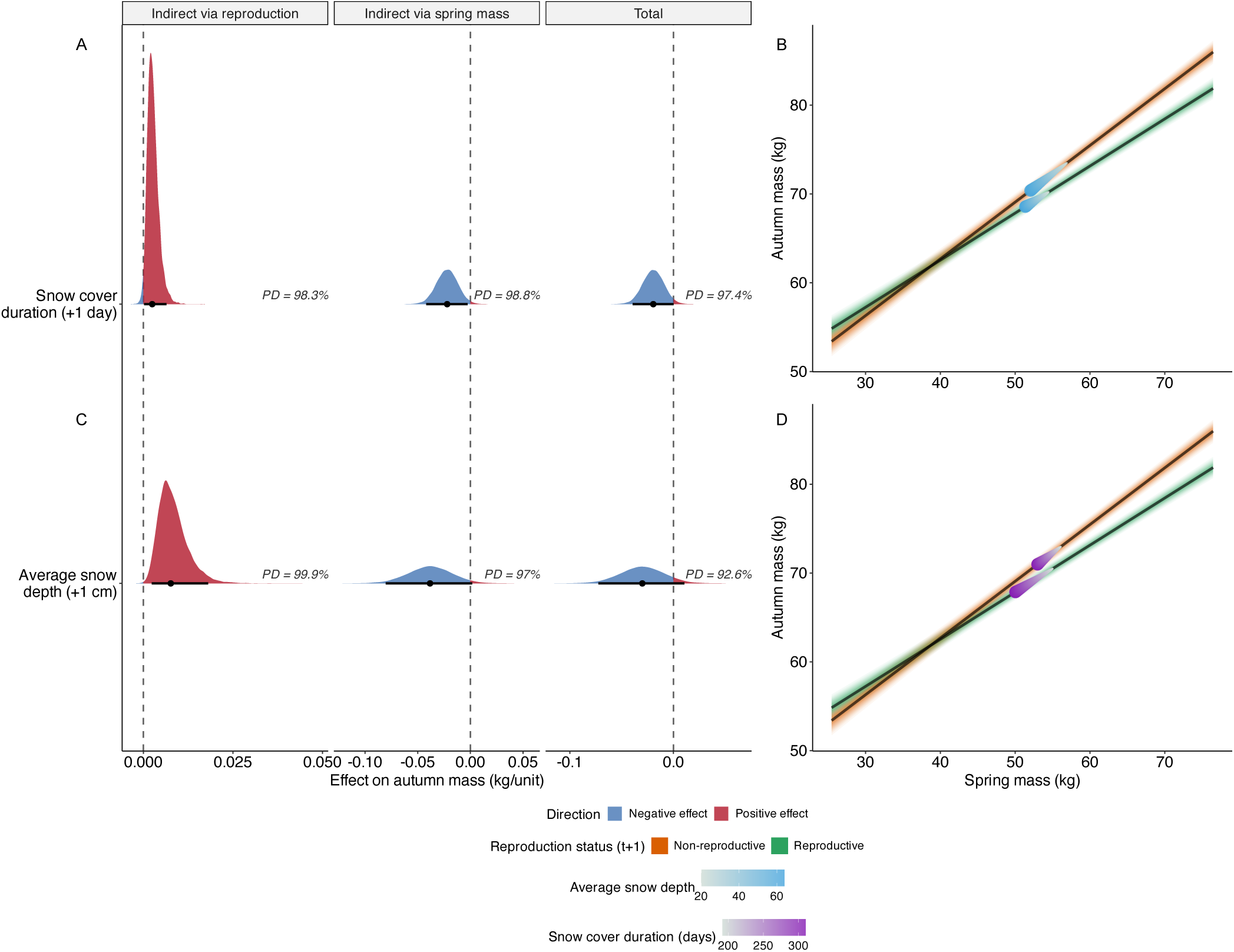
Decomposition of carry-over effects of snow conditions on subsequent autumn body mass in adult female bighorn sheep (2 yr+ at *t+1*). The analysis decomposes the total effect of (A) snow cover duration and (C) average snow depth into two distinct pathways: Indirect via reproduction (representing the mass saved by a reduction in reproduction probability) and Indirect via spring mass (representing somatic costs of snow). The left panels show posterior distributions of the standardized effects. Density ridges are colored by direction: red indicates a positive effect (buffering) and blue a negative effect (cost). Black points indicate the posterior median, and horizontal bars show 95% credible intervals. Vertical dashed lines denote no effect (0). Values report the probability of direction (PD). The right panels show the visualization of indirect effect magnitudes following Bush-Beaupre et al. (2026). The black regression lines (B, D) depict the relationship between spring mass and autumn mass. The superimposed colored segments (blue: snow depth; purple: snow cover duration) illustrate the indirect pathway. Reading the segments along their color gradient illustrates the indirect effect: as snow conditions become more severe, the impact induced on spring mass (the shift along the x-axis) subsequently dictates a proportional change in autumn mass (shifting down on the y-axis).

Population density strongly modulated carry-over effects in males. We found an effect of snowpack density on next autumn body mass only at high population density: an increase in snow density of 1 kg/m³ resulted in a decrease of 0.04 kg in autumn body mass (95% CI [-0.14; 0.07]) (Figure 7D). At high density, males experiencing the densest snow were ∼3 kg (-3.18% of mean mass) lighter the following autumn than those experiencing soft snow (Figure 7D). In contrast, at low population density, this carry-over effect was positive (0.09 kg per kg/m³, 95% CI [0.02; 0.18]) (Figure 7C), leading to a mass gain of ∼7 kg (7.97% of mean mass) across the range of snow density (Figure 7D). Similarly, the carry-over effect of average snow depth was dependent on population density. At low and mean population densities, an increase of 1 cm in snow depth resulted in a strong mass loss of 0.32 kg (95% CI [-0.47; -0.16]) and 0.25 kg (95% CI [-0.36; -0.13]), respectively. However, at high population density, this negative effect weakened (-0.17 kg/cm, 95% CI [-0.36; 0.02]). In contrast, snow cover duration induced a carry-over effect that remained independent of population density, with a decrease of 0.06 kg per day (95% CI [-0.12; -0.01], Figure 7E), leading to a decrease of ∼7.2 kg (-8.34% of mean mass) (Figure 7F) in autumn mass between the shortest and the longest snow conditions.

**Figure 7:**
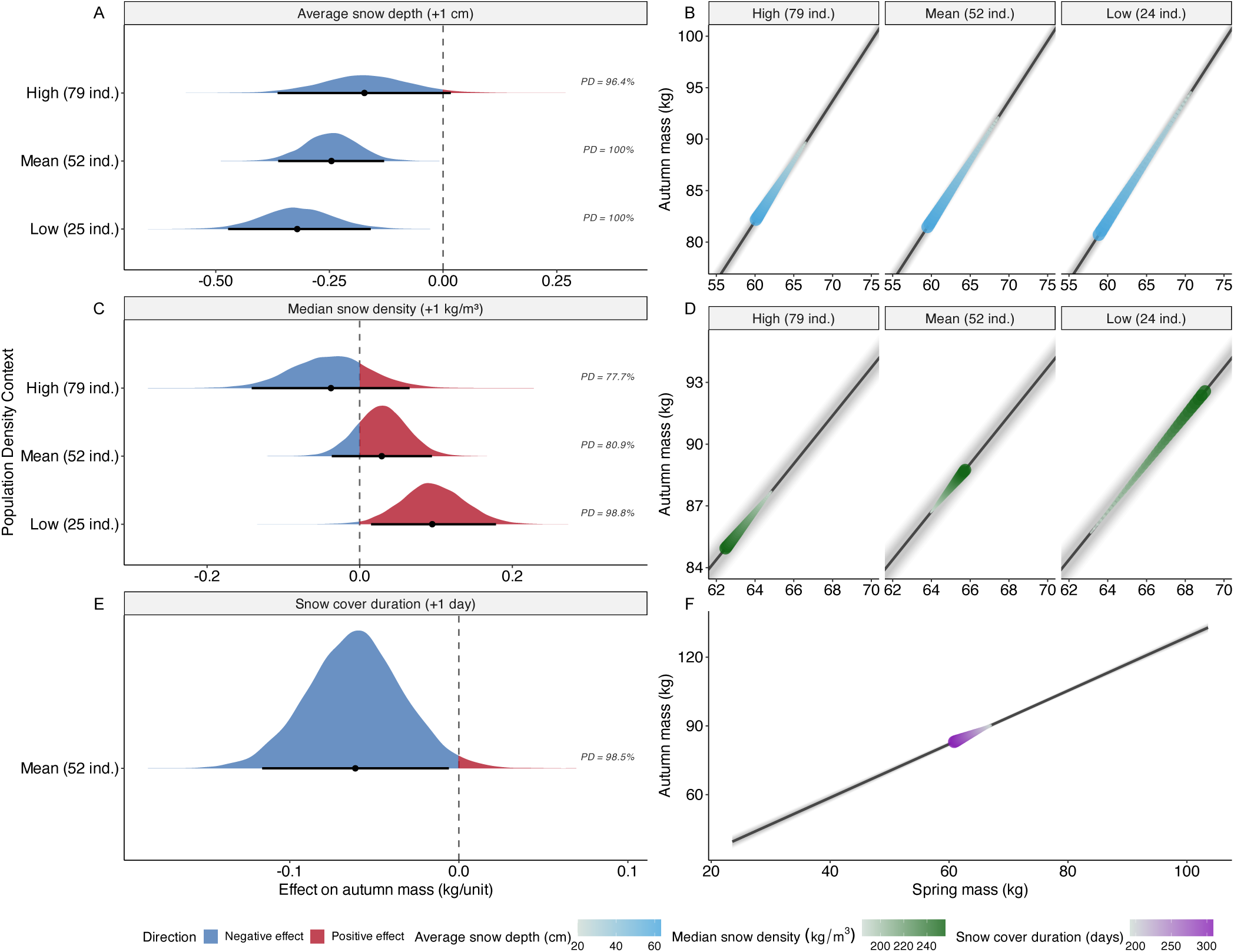
Context-dependent carry-over effects of snow conditions on subsequent autumn body mass in adult male bighorn sheep (2 yr+ at *t+1*). Left panels show the posterior distributions of the effects of (A) average snow depth, (C) median snow density, and (E) snow cover duration on autumn body mass. For snow depth (B) and density (C), estimates are shown across three population densities: low (25 ind.), mean (52 ind.), and high (79 ind.), revealing a reversal in effect direction. Density ridges are colored by direction (red = positive; blue = negative). Black points indicate the posterior median with 95% credible intervals. Values report the Probability of Direction (PD). Right panels show the visualization of carry-over effect magnitudes following Bush-Beaupre et al. (2026). The regression lines (B, D, F) show the relationship between spring mass and autumn mass. The superimposed colored segments (blue: snow depth; green: median snow density; purple: snow cover duration) illustrate the indirect pathway. Reading the segments along their color gradient illustrates the indirect effect: as snow conditions become more severe, the impact induced on spring mass (the shift along the x-axis) subsequently dictates a proportional change in autumn mass (the resulting shift along the y-axis).

## Discussion

Our study demonstrates that snow conditions are a critical environmental constraint on bighorn sheep performance, yet the magnitude and mechanisms of these effects vary markedly with demographic group and population density. While deep snow and long duration of snow cover consistently lowered spring body mass across all sex-age groups, the influence of snow density was complex and context-dependent, particularly for males. Snow impacts include carry-over effects on body mass in the subsequent autumn for all groups. However, the pathways differed: females appeared to partially buffer somatic costs through reproductive trade-offs, whereas yearlings faced unmitigated negative effects of difficult snow conditions.

Harsh snow conditions acted as a broad limiting factor. Both snow depth and duration were consistently associated with reduced spring mass, likely by reducing dry matter intake and digestibility (Goodson et al. 1991), while simultaneously increasing locomotion costs. These effects were particularly pronounced in lambs, which have lower energy reserves than adults (Parker et al., 1984). The heightened sensitivity of lambs likely reflects mechanical constraints linked to smaller body size; shorter legs increase the energetic cost of movement in deep snow (Parker et al., 1984), making lambs especially vulnerable to heavy snow accumulation.

For females, our findings align with Festa-Bianchet (1998), who documented greater overwinter mass loss in reproductive individuals. Prolonged snow cover appears to compound forage limitations with the high energetic demands of gestation. The negative relationships between snow duration and depth with reproductive probability suggest a physiological buffering mechanism: under harsh conditions, females may fail to conceive or may terminate gestation to preserve somatic condition. Similar responses have been documented in other ungulates (Dimac-Stohl et al., 2018; Parker et al., 2009). While Mysterud & Østbye (2006) did not specifically consider reproductive rates in roe deer (*Capreolus capreolus*), their findings highlight that severe winters drive broad-scale declines in both autumn body mass and population growth, likely reflecting the cumulative impact of reduced individual performance and lower survival.

In adult males, snow density appeared to have a density-dependent effect on spring body mass. At low population density, increasing snow density was associated with higher spring body mass, whereas this relationship weakened at intermediate density and trended negative at high population density. This reversal in direction suggests that the effect of snow density on male body condition cannot be interpreted independently of density-dependent processes. This finding is coherent with a study on Soay sheep (*Ovis aries*), where the impact of harsh winter weather similarly increased with population density (Coulson et al., 2001). At low population density, denser snow may facilitate movement by providing greater load-bearing capacity, thereby reducing locomotion costs compared to softer snow (Telfer & Kelsall, 1979). Furthermore, at low density, individuals may benefit from this increased mobility to access forage with minimal competition. At higher population density, increased competition for limited snow-free patches (such as wind-blown ridges or steep slopes) may offset the locomotory advantages provided by denser snow. While denser snow might also increase the cost of cratering for forage (Skogland, 1978), the primary mechanism at high density appears to be the saturation of optimal wintering areas, where social interactions and competition for restricted resources overwhelm the benefits of improved snow support. Under these conditions, denser snow ultimately results in reduced spring body mass. While more uncertain, we also found evidence that the negative impact of average snow depth on male spring mass weakened and trended towards zero at high population densities. We hypothesize that this buffering effect could be explained by collective behaviors and snowpack alteration. A larger number of individuals moving in single file may help to pack the snow on the trail, which significantly reduces the energetic costs of locomotion through deep snow (Telfer & Kelsall, 1984). These effects also extend beyond the snow season, suggesting delayed carry-over consequences for body mass later in the year.

Our results provide strong evidence that harsh snow conditions generate persistent carry-over effects on body mass in the following autumn (Harrison et al., 2011; O’connor et al., 2014). Therefore, easy access to food from June to mid-September cannot fully compensate for several months of food shortages. Therefore, easy access to food from June to mid-September cannot fully compensate for several months of food shortages. However, the mechanisms driving these delayed effects differ by group, revealing a higher sensitivity of males to snow conditions.

In adult females, the negative carry-over impact of severe snow conditions was partially offset by a positive effect mediated through reproduction. Harsh snow conditions reduced reproductive probability, thereby saving the female the energetic costs of late gestation and lactation (Festa-Bianchet, 1998). Consequently, while the total effect of snow severity on autumn mass remained negative, it was modulated by lower allocation to reproduction. This highlights reproductive modulation as a key strategy allowing females to prioritize somatic maintenance under difficult conditions (Martin & Festa-Bianchet, 2010; Therrien et al., 2007).

In contrast, lambs that survived their first winter showed strong, unmitigated carry-over effects as yearlings on autumn body mass. Increased snow depth and duration led to substantial reductions in yearling autumn mass. Male yearlings exhibited the highest sensitivity to snow depth of any sex-age group, consistent with the hypothesis that higher growth demands increase sensitivity to environmental stress in the faster-growing sex (Clutton-Brock et al., 1985). Lacking the ability to reduce allocation to reproduction, yearlings absorb winter energy deficits directly as reduced growth. Importantly, our results likely underestimate these impacts. We could only record the mass of individuals that survived the winter, as those that suffered the most severe mass loss likely died before spring. The pronounced sensitivity of lambs (Gaillard et al., 2000) likely reflects limited energy reserves, small body size, and higher locomotion costs under deep or persistent snow cover (Telfer & Kelsall, 1979). Severe snow conditions during early life could have long-term consequences for cohort quality and survival (Pigeon & Pelletier, 2018).

Consistent with the observations in yearlings, adult males also displayed significant sensitivity to snow conditions. In this group, carry-over effects mirrored the context-dependence observed in spring. Snow density showed a pronounced density-dependent carry-over effect: beneficial at low density but likely detrimental at high density. We hypothesize that the persistence of snow effects reflects insufficient compensatory summer growth under high-density conditions. Since spring mass determines the baseline for summer mass gain, males exiting the snow season with a heavy energetic deficit, imposed by the combination of deep snow, dense snow and high competition, struggle to recover. While ungulates typically exhibit compensatory growth following resource restriction (Rughetti & Festa-Bianchet, 2011), high population density during summer likely constrains nutrient intake, preventing males from recovering the mass lost through dense snow (Leblanc et al., 2001). The persistent effect of the previous snow conditions on autumn body mass suggests a cumulative energetic debt that transcends a single season. Because body mass is a primary determinant of overwinter survival and reproductive potential in bighorn sheep (Festa-Bianchet et al., 1997), individuals entering the subsequent snow season with a mass deficit face a compounded risk. A reduced mass at the onset of snow season limits the ability to buffer environmental constraints, thereby increasing the probability of mortality or reproductive failure in year *t+2*. The demographic effect of severe snow conditions may thus be felt for several years through time-lagged effects on individual fitness.

## Conclusion

The effects of harsh snow conditions on sheep cannot be inferred from snow depth alone. Snow persistence, physical structure, and population density jointly determine how individuals experience winter. While females may mitigate long-term effects of harsh conditions through reproductive adjustments, juveniles or males do not have that flexibility. Males also face density-dependent constraints. Crucially, the reduction in autumn body mass driven by snow conditions likely has cascading phenotypic consequences. Because body mass is a primary determinant of overwinter survival and reproductive potential, individuals entering the subsequent snow season with a mass deficit may face lower survival and reproduction probability. As climate change alters snowpack properties and persistence, understanding these mechanistic drivers will be critical for predicting the dynamics of large herbivore populations in seasonal environments.

## Author contributions

K.C., F.P., and M.F-B. conceptualized the study and performed data collection. K.C. performed the statistical analyses, interpreted the results, and drafted the manuscript. F.P., M.F-B., and A.L. contributed to the writing and critical revision of the manuscript. M.F-B. and F.P. are responsible for the long-term research program at Ram Mountain.

## Acknowledgments

We are grateful to the many students, colleagues, and research assistants who have participated in data collection since 1971, with special thanks to J. Jorgenson. We also thank C. Feder and A. Hubbs for their continued support of the Ram Mountain research program.

## Data availability statement

Data and code are available in an Open Science Framework repository (https://doi.org/10.17605/OSF.IO/F5M9V)

