## Appendix S1 for "Harsh snow conditions reduce body mass and reproductive success of an alpine ungulate, inducing carry-over effects"

**Table S1**: List of prediction tested for Spring body mass and reproductive probability, along with the corresponding population group for bighorn sheep on Ram Mountain, Alberta, Canada

| Prediction | Population group |
| --- | --- |
| **1)** Spring body mass was negatively associated with winter conditions and positively associated with autumn body mass; in turn, autumn body mass was positively related to body mass in the following autumn. | Both |
| **2)** Spring body mass was negatively associated with winter conditions through an interaction with age and positively associated with autumn body mass, which was itself positively related to body mass in the subsequent autumn. | Males and females 1 year+ |
| **3)** Spring body mass was negatively associated with winter conditions, with a stronger effect under high population density, and positively associated with autumn body mass; autumn body mass was positively related to body mass in the following autumn. | Both |
| **4)** Spring body mass was negatively associated with winter conditions, with a stronger effect in pregnant females, and positively associated with autumn body mass; autumn body mass was positively related to body mass in the subsequent autumn. | Females 1 year+ |

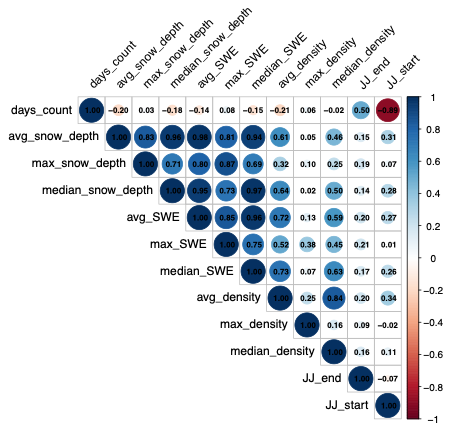

**Figure S1**: Correlation matrix between snow characteristic variables (duration of snow cover (days_count), mean, maximum, median of snow cover depth(*_snow_depth), mean, maximum, median of snow water equivalent (*_SWE) and mean, maximum and median of snow cover density (*_density)), end (JJ_end) and start date (JJ_start).

**Table S2**: Comparison of age modeling for females and males aged 1 year and over. For each population group, three types of modeling were tested: linear modeling, quadratic modeling and nonlinear modeling with a natural 3rd degree spline. Presented for the three response variables tested: spring mass, next autumn mass and reproduction status. Model comparison is based on ELPD. The model with the lowest ELPD was selected as the best.

| **Response**  **variable** | **Population group** | **Model type** | **ELPD** | **∆ ELPD** |
| --- | --- | --- | --- | --- |
| Spring mass | Female 1 yr+ | Linear | -5233.5 | -648.5 |
|  | Female 1 yr+ | Quadratic | -4788.3 | -203.3 |
|  | **Female 1 yr+** | **Spline 3^rd^ degree** | **-4585** | **0.00** |
|  | Male 1 yr+ | Linear | -1745.1 | -348.9 |
|  | Male 1 yr+ | Quadratic | -1429.7 | -33.5 |
|  | **Male 1 yr+** | **Spline 3^rd^ degree** | **-1396.2** | **0.00** |
| Next autumn mass | Female 1 yr+ | Linear | -5043.3 | -774.5 |
|  | Female 1 yr+ | Quadratic | -4477.3 | -208.5 |
|  | **Female 1 yr+** | **Spline 3^rd^ degree** | **-4268.8** | **0.00** |
|  | Male 1 yr+ | Linear | **-1799.4** | - 306.7 |
|  | Male 1 yr+ | Quadratic | -1513.6 | -21 |
|  | **Male 1 yr+** | **Spline 3^rd^ degree** | **-1492.7** | **0.00** |
| Reproduction status | Female 1 yr+ | Linear | -729.0 | -220.6 |
|  | Female 1 yr+ | Quadratic | - 544.7 | -36.3 |
|  | **Female 1 yr+** | **Spline 3^rd^ degree** | **-** **508.4** | **0.00** |

**Table S3**: The structural equation modelling was performed separately for weaned lambs, adult females, and adult males. The table details the fixed effects structure for each response variable (Spring mass, Next autumn mass, and Reproduction probability) within each model. Snow variables represent the set of three winter covariates: average snow depth, snow cover duration, and median snow density. All models included random intercepts for Year and Individual ID for adults and only Year for weaned lambs to account for repeated measures and/or inter-annual variability.

| **Groupe** | **Model** | **Response variable** | **Fixed effect** |
| --- | --- | --- | --- |
| LAMBS | Base model | Spring mass | Sex * (Autumn mass + Population density + Snow variables) |
|  |  | Next autumn mass | Sex * (Spring mass + Next population density ) |
|  | Population density in interaction with snow variables | Spring mass | Sex * Automn mass + Sex * Population density * Snow variables |
|  |  | Next autumn mass | Sex * (Spring mass + Next population density ) |
| FEMALES | Base model | Reproduction probability | Age (spline) + Autumn Mass + Population density + Snow variables |
|  |  | Spring mass | Autumn Mass + Age + Population density + Reproductive status + Snow variables |
|  |  | Next autumn mass | Reproductive status * Spring mass + Age next year (spline) + Next population density |
|  | Age interaction with snow variables | Reproduction probability | Age (spline) + Autumn Mass + Population density + Snow variables |
|  |  | Spring mass | ... + Age * Snow variables |
|  |  | Next autumn mass | Reproductive status * Spring mass + Age next year (spline) + Next population density |
|  | Population density in interaction with snow variables | Reproduction probability | Age (spline) + Autumn Mass + Population density + Snow variables |
|  |  | Spring mass | ... + Population density * Snow variables |
|  |  | Next autumn mass | Reproductive status * Spring mass + Age next year (spline) + Next population density |
|  | Reproduction status in interaction with snow variables | Reproduction probability | Age (spline) + Autumn Mass + Population density + Snow variables |
|  |  | Spring mass | ... + Reproductive status * Snow variables |
|  |  | Next autumn mass | Reproductive status * Spring mass + Age next year (spline) + Next population density |
| MALES | Base model | Spring mass | Autumn Mass + Age (spline) + Population density + Snow variables |
|  |  | Next autumn mass | Spring mass + Age Next (spline) + Next population density |
|  | Age interaction with snow variables | Spring mass | ... + Age (spline) * Snow variables |
|  |  | Next autumn mass | Spring mass + Age Next (spline) + Next population density |
|  | Population density in interaction with snow variables | Spring mass | ... + Population density * Snow variables |
|  |  | Next autumn mass | Spring mass + Age Next (spline)+ Next population density |

**Table S4**: Power-scaling sensitivity analysis for Male, Female, and Lamb models. Sensitivity was assessed using the Cumulative Jensen–Shannon distance via the *priorsense* package. Prior: Quantifies the change in the posterior distribution when the prior is perturbed; values near 0 indicate that the prior does not unduly constrain the posterior. Likelihood: Quantifies the influence of the data; values > 1.0 indicate that the data are highly informative and dominate the posterior distribution. Diagnosis: Checks for prior–data conflicts; a hyphen (–) indicates no significant conflict was detected.

| **Model** | **Response variable** | **Parameter** | **Prior** | **Likelihood** | **Diagnosis** |
| --- | --- | --- | --- | --- | --- |
| MALES | *Spring Mass* | Intercept | 0.000 | 2.740 | – |
|  |  | Autumn Mass (t-1) | 0.000 | 3.158 | – |
|  |  | Age Spline 1 | 0.000 | 2.989 | – |
|  |  | Age Spline 2 | 0.000 | 3.009 | – |
|  |  | Age Spline 3 | 0.000 | 2.837 | – |
|  |  | Pop Density | 0.000 | 0.255 | – |
|  |  | Avg Snow Depth | 0.000 | 0.113 | – |
|  |  | Snow Cover Duration | 0.000 | 0.125 | – |
|  |  | Median Snow Density | 0.000 | 0.137 | – |
|  |  | Pop Density × Avg Snow Depth | 0.000 | 0.072 | – |
|  |  | Pop Density × Snow Cover Duration | 0.000 | 0.074 | – |
|  |  | Pop Density × Median Snow Density | 0.000 | 0.148 | – |
|  |  | SD ID (Intercept) | 0.000 | 3.521 | – |
|  |  | SD Year (Intercept) | 0.001 | 0.544 | – |
|  |  | Sigma | 0.000 | 3.759 | – |
|  |  | Intercept† | 0.000 | 0.192 | – |
|  | *Next Autumn Mass* | Intercept | 0.000 | 0.577 | – |
|  |  | Spring Mass | 0.000 | 1.004 | – |
|  |  | Age Spline (Next) 1 | 0.000 | 0.702 | – |
|  |  | Age Spline (Next) 2 | 0.000 | 0.927 | – |
|  |  | Age Spline (Next) 3 | 0.000 | 0.843 | – |
|  |  | Next Pop Density | 0.000 | 0.070 | – |
|  |  | SD ID (Intercept) | 0.000 | 2.444 | – |
|  |  | SD Year Next (Intercept) | 0.001 | 0.091 | – |
|  |  | Sigma | 0.000 | 2.170 | – |
|  |  | Intercept† | 0.000 | 0.136 | – |
| FEMALES | *Reproduction status* | Intercept | 0.007 | 0.075 | – |
|  |  | Age Spline 1 | 0.004 | 0.086 | – |
|  |  | Age Spline 2 | 0.004 | 0.063 | – |
|  |  | Age Spline 3 | 0.003 | 0.358 | – |
|  |  | Autumn Mass (t-1) | 0.008 | 0.609 | – |
|  |  | Pop Density | 0.004 | 0.147 | – |
|  |  | Avg Snow Depth | 0.005 | 0.153 | – |
|  |  | Snow Cover Duration | 0.005 | 0.159 | – |
|  |  | Median Snow Density | 0.002 | 0.133 | – |
|  |  | SD ID (Intercept) | 0.017 | 1.130 | – |
|  |  | SD Year (Intercept) | 0.022 | 0.639 | – |
|  |  | Intercept† | 0.021 | 0.308 | – |
|  | *Spring Mass* | Intercept | 0.002 | 1.265 | – |
|  |  | Autumn Mass (t-1) | 0.001 | 1.679 | – |
|  |  | Age Spline 1 | 0.000 | 1.212 | – |
|  |  | Age Spline 2 | 0.001 | 1.454 | – |
|  |  | Age Spline 3 | 0.000 | 0.625 | – |
|  |  | Pop Density | 0.001 | 0.261 | – |
|  |  | Repro Status | 0.000 | 0.167 | – |
|  |  | Avg Snow Depth | 0.000 | 0.112 | – |
|  |  | Snow Cover Duration | 0.001 | 0.074 | – |
|  |  | Median Snow Density | 0.000 | 0.124 | – |
|  |  | Repro Status × Avg Snow Depth | 0.001 | 0.171 | – |
|  |  | Repro Status × Snow Cover Duration | 0.000 | 0.118 | – |
|  |  | Repro Status × Median Snow Density | 0.000 | 0.145 | – |
|  |  | SD ID (Intercept) | 0.001 | 1.726 | – |
|  |  | SD Year (Intercept) | 0.004 | 0.075 | – |
|  |  | Sigma | 0.001 | 1.329 | – |
|  |  | Intercept† | 0.002 | 0.170 | – |
|  | *Next Autumn Mass* | Intercept | 0.002 | 0.564 | – |
|  |  | Repro Status | 0.000 | 0.334 | – |
|  |  | Spring Mass | 0.000 | 1.181 | – |
|  |  | Age Spline (Next) 1 | 0.000 | 0.868 | – |
|  |  | Age Spline (Next) 2 | 0.000 | 0.929 | – |
|  |  | Age Spline (Next) 3 | 0.001 | 0.326 | – |
|  |  | Next Pop Density | 0.001 | 0.098 | – |
|  |  | Repro Status × Spring Mass | 0.000 | 0.222 | – |
|  |  | SD ID (Intercept) | 0.000 | 1.074 | – |
|  |  | SD Year Next (Intercept) | 0.003 | 0.149 | – |
|  |  | Sigma | 0.001 | 1.355 | – |
|  |  | Intercept† | 0.002 | 0.046 | – |
| LAMBS | *Spring Mass* | Intercept | 0.001 | 0.029 | – |
|  |  | Autumn Mass (t-1) | 0.000 | 0.094 | – |
|  |  | Male | 0.000 | 0.097 | – |
|  |  | Pop Density | 0.000 | 0.024 | – |
|  |  | Avg Snow Depth | 0.000 | 0.031 | – |
|  |  | Snow Cover Duration | 0.000 | 0.016 | – |
|  |  | Median Snow Density | 0.000 | 0.047 | – |
|  |  | Male × Autumn Mass (t-1) | 0.000 | 0.097 | – |
|  |  | Male × Pop Density | 0.000 | 0.100 | – |
|  |  | Male × Avg Snow Depth | 0.000 | 0.097 | – |
|  |  | Male × Snow Cover Duration | 0.000 | 0.094 | – |
|  |  | Male × Median Snow Density | 0.000 | 0.097 | – |
|  |  | SD Year (Intercept) | 0.005 | 0.136 | – |
|  |  | Sigma | 0.001 | 0.384 | – |
|  |  | Intercept† | 0.001 | 0.040 | – |
|  | *Next Autumn Mass* | Intercept | 0.001 | 0.059 | – |
|  |  | Male | 0.000 | 0.112 | – |
|  |  | Spring Mass | 0.000 | 0.194 | – |
|  |  | Next Pop Density | 0.000 | 0.033 | – |
|  |  | Male × Spring Mass | 0.000 | 0.114 | – |
|  |  | Male × Next Pop Density | 0.000 | 0.122 | – |
|  |  | SD Year Next (Intercept) | 0.003 | 0.213 | – |
|  |  | Sigma | 0.001 | 0.333 | – |
|  |  | Intercept† | 0.001 | 0.030 | – |

**†** Intercept (marginal): the uncentered population-level intercept as reported by *priorsense* for multivariate brms models; distinct from the design-matrix intercept shown in the main parameter list.

**Table S5**: Comparison of models for each prediction (prediction number from Table S1) for each corresponding population group. Selection criteria include ELPD and ΔELPD.

| **Population group** | **Prediction** | **ELPD** | **∆ELPD** |
| --- | --- | --- | --- |
| **Lamb** | **1** | -1775.3 | **-0.00** |
| Lamb | 3 | -1777.3 | -2 |
| Female 1 yr+ | 1 | -8861.3 | -0.6 |
| Female 1 yr+ | 2 | -8861.4 | -0.7 |
| Female 1 yr+ | 3 | -8862.0 | -1.3 |
| **Female 1 yr+** | **4** | **-8860.7** | **0** |
| Male 1 yr+ | 1 | -2670.2 | -5.3 |
| Male 1 yr+ | 2 | -2675.8 | -10.9 |
| **Male 1 yr+** | **3** | **-2576.2** | **0.00** |

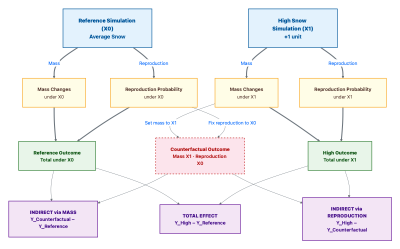

**Figure S2**: Comparing a Reference Simulation (X_0_, e.g., average snow conditions) against a High Snow Simulation (X_1_, e.g., +1 unit). To disentangle the mechanisms, a Counterfactual Outcome (dashed red box) is simulated representing a hypothetical scenario where individuals experience the mass changes associated with high snow (X_1_) while maintaining the reproduction probability of the reference conditions (X_0_). This decomposition partitions the total effect into two pathways (purple boxes): (1) Indirect Effect via Mass (calculated as Y _Counterfactual_ – Y _Reference_): captures the cost of snow on body mass independent of changes in reproductive status. (2) Indirect Effect via Reproduction (calculated as Y _High_ – Y _Counterfactual_): captures the change in body mass resulting solely from the snow-induced shift in reproductive success.
