## Appendix S2 for "Harsh snow conditions reduce body mass and reproductive success of an alpine ungulate, inducing carry-over effects"

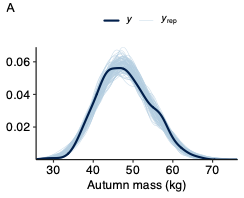

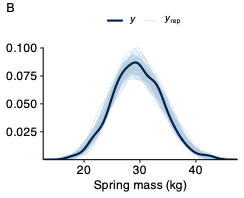

**Figure S1**: Posterior predictive check for autumn mass (A) and spring mass (B) for the lambs model. The thick dark blue line represents the kernel density estimate of the observed data (*y*). The thinner light blue lines represent kernel density estimates of the 100 replicated datasets (*y_rep_*) drawn from the posterior predictive distribution. The strong overlap between the observed data and the simulated replications indicates that the model adequately captures the distribution of the observed data.

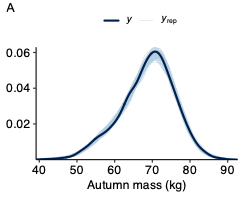

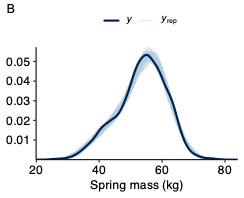

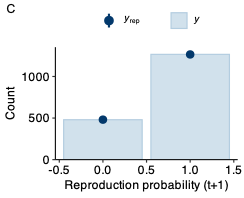

**Figure S2**: Posterior predictive check for autumn mass (A), spring mass (B) and the reproduction probability (C) for the adult females model. The thick dark blue line represents the kernel density estimate of the observed data (*y*). The thinner light blue lines represent kernel density estimates of the 100 replicated datasets (*y_rep_*) drawn from the posterior predictive distribution. The strong overlap between the observed data and the simulated replications indicates that the model adequately captures the distribution of the observed data.

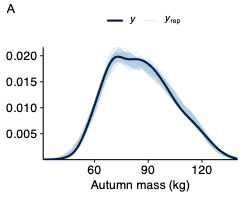

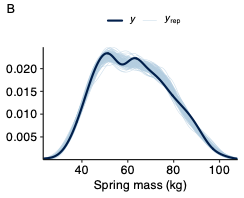

**Figure S3**: Posterior predictive check for autumn mass (A) and spring mass (B) for the adult males model. The thick dark blue line represents the kernel density estimate of the observed data (*y*). The thinner light blue lines represent kernel density estimates of the 100 replicated datasets (*y_rep_*) drawn from the posterior predictive distribution. The strong overlap between the observed data and the simulated replications indicates that the model adequately captures the distribution of the observed data.
