## Appendix S3 for "Harsh snow conditions reduce body mass and reproductive success of an alpine ungulate, inducing carry-over effects"

**Table S1**: Posterior estimates from the Bayesian structural equation models assessing the effect of age, body mass, reproductive status, population density and snow conditions on spring body mass on three sex-age groups of bighorn sheep on Ram Mountain, Alberta. For each response variable, posterior mean estimates, standard errors (Est.Error) and 95% credible intervals (l-95% CI to u-95% CI) are presented. Convergence diagnostics include the R-hat statistic (Rhat) and effective sample sizes for the bulk and tail of the posterior distributions (Bulk_ESS and Tail_ESS. respectively). All predictors were standardized prior to model fitting.

| Population group | Response | Intercept/Fixed effect | Estimate | Est.Error | l-95% CI | u-95% CI | Rhat | Bulk_ESS | Tail_ESS |
| --- | --- | --- | --- | --- | --- | --- | --- | --- | --- |
| Lambs | Spring mass | Intercept | 28.821 | 0.310 | 28.216 | 29.427 | 1.0 | 5245 | 8076 |
|  | Next autumn mass | Intercept | 46.313 | 0.369 | 45.588 | 47.047 | 1.0 | 6027 | 8939 |
|  | Spring mass | Sex (Male) | 1.464 | 0.240 | 0.994 | 1.937 | 1.0 | 16831 | 12433 |
|  |  | Autumn mass | 2.761 | 0.192 | 2.391 | 3.135 | 1.0 | 10067 | 11302 |
|  |  | Population density | -0.045 | 0.296 | -0.626 | 0.538 | 1.0 | 5265 | 8136 |
|  |  | Average snow depth | -0.865 | 0.358 | -1.566 | -0.160 | 1.0 | 4834 | 8084 |
|  |  | Snow cover duration | -0.800 | 0.322 | -1.433 | -0.172 | 1.0 | 5423 | 8395 |
|  |  | Median snow density | -0.026 | 0.359 | -0.727 | 0.690 | 1.0 | 4754 | 7642 |
|  |  | Autumn mass : Sex (Male) | 0.277 | 0.241 | -0.194 | 0.744 | 1.0 | 11347 | 11034 |
|  |  | Population density : Sex (Male) | -0.619 | 0.226 | -1.065 | -0.173 | 1.0 | 14896 | 11960 |
|  |  | Average snow depth : Sex (Male) | -0.461 | 0.279 | -1.002 | 0.084 | 1.0 | 11298 | 12993 |
|  |  | Snow cover duration : Sex (Male) | -0.019 | 0.240 | -0.488 | 0.454 | 1.0 | 14265 | 12795 |
|  |  | Median snow density : Sex (Male) | 0.095 | 0.270 | -0.433 | 0.629 | 1.0 | 11154 | 11787 |
|  | Next autumn mass | Sex (Male) | 2.631 | 0.351 | 1.948 | 3.316 | 1.0 | 12662 | 12313 |
|  |  | Spring mass | 4.753 | 0.332 | 4.097 | 5.405 | 1.0 | 6981 | 9399 |
|  |  | Next population density | -1.129 | 0.343 | -1.798 | -0.463 | 1.0 | 6004 | 8716 |
|  |  | Spring mass : Sex (Male) | 0.912 | 0.347 | 0.234 | 1.586 | 1.0 | 12066 | 12016 |
|  |  | Next population density : Sex (Male) | 0.130 | 0.310 | -0.486 | 0.735 | 1.0 | 15178 | 11837 |
| Females 1 yr+ | Reproduction | Intercept | -0.895 | 0.632 | -2.159 | 0.317 | 1.0 | 7524 | 9683 |
|  | Spring mass | Intercept | 53.251 | 0.734 | 51.775 | 54.677 | 1.0 | 3102 | 6239 |
|  | Next autumn mass | Intercept | 65.489 | 0.492 | 64.530 | 66.447 | 1.0 | 2318 | 5117 |
|  | Reproduction | Age. 1 | 2.985 | 0.785 | 1.479 | 4.564 | 1.0 | 11813 | 11869 |
|  | Reproduction | Age. 2 | 5.028 | 1.507 | 2.155 | 8.060 | 1.0 | 8003 | 10399 |
|  | Reproduction | Age. 3 | -2.964 | 1.144 | -5.234 | -0.754 | 1.0 | 9866 | 10594 |
|  | Reproduction | Autumn mass | 2.141 | 0.324 | 1.531 | 2.793 | 1.0 | 7520 | 10405 |
|  | Reproduction | Population density | -0.804 | 0.329 | -1.469 | -0.181 | 1.0 | 5555 | 8019 |
|  | Reproduction | Average snow depth | -0.978 | 0.315 | -1.610 | -0.376 | 1.0 | 5439 | 8450 |
|  | Reproduction | Snow cover duration | -0.630 | 0.293 | -1.219 | -0.074 | 1.0 | 6548 | 8482 |
|  | Reproduction | Median snow density | -0.023 | 0.308 | -0.639 | 0.593 | 1.0 | 6026 | 8684 |
|  | Spring mass | Autumn mass | 6.321 | 0.299 | 5.741 | 6.907 | 1.0 | 3576 | 7174 |
|  | Spring mass | Age. 1 | 2.038 | 0.668 | 0.737 | 3.333 | 1.0 | 5651 | 9696 |
|  | Spring mass | Age. 2 | 3.188 | 1.450 | 0.367 | 6.036 | 1.0 | 4042 | 7428 |
|  | Spring mass | Age. 3 | 0.897 | 0.928 | -0.952 | 2.684 | 1.0 | 9482 | 11078 |
|  | Spring mass | Population density | 0.010 | 0.463 | -0.896 | 0.932 | 1.0 | 2998 | 4781 |
|  | Spring mass | Reproduction status | -1.760 | 0.249 | -2.251 | -1.274 | 1.0 | 18894 | 12678 |
|  | Spring mass | Average snow depth | -1.258 | 0.466 | -2.179 | -0.345 | 1.0 | 3546 | 6254 |
|  | Spring mass | Snow cover duration | -0.592 | 0.434 | -1.446 | 0.255 | 1.0 | 3895 | 6094 |
|  | Spring mass | Median snow density | 0.368 | 0.464 | -0.552 | 1.262 | 1.0 | 3329 | 5784 |
|  | Spring mass | Reproduction status : Average snow depth | 0.454 | 0.200 | 0.064 | 0.852 | 1.0 | 16279 | 13008 |
|  | Spring mass | Reproduction status : Snow cover duration | -0.352 | 0.169 | -0.686 | -0.020 | 1.0 | 21663 | 12451 |
|  | Spring mass | Reproduction status : Median snow density | -0.002 | 0.203 | -0.401 | 0.400 | 1.0 | 16680 | 12881 |
|  | Next autumn mass | Reproduction status | -1.608 | 0.185 | -1.967 | -1.247 | 1.0 | 11572 | 11356 |
|  | Next autumn mass | Spring mass | 5.054 | 0.184 | 4.692 | 5.419 | 1.0 | 4875 | 7723 |
|  | Next autumn mass | Age_t+1. 1 | 5.825 | 0.421 | 5.000 | 6.646 | 1.0 | 7664 | 10366 |
|  | Next autumn mass | Age_t+1. 2 | 9.430 | 0.839 | 7.805 | 11.082 | 1.0 | 4526 | 6729 |
|  | Next autumn mass | Age_t+1. 3 | 0.558 | 0.663 | -0.745 | 1.854 | 1.0 | 7728 | 10115 |
|  | Next autumn mass | Next population density | -0.888 | 0.399 | -1.680 | -0.113 | 1.0 | 2166 | 4134 |
|  | Next autumn mass | Spring mass : Reproduction status | -0.860 | 0.153 | -1.161 | -0.557 | 1.0 | 12223 | 11781 |
| Males 1 yr+ | Spring mass | Intercept | 60.651 | 1.176 | 58.325 | 62.979 | 1.0 | 2253 | 4277 |
|  | Next autumn mass | Intercept | 85.694 | 0.751 | 84.227 | 87.170 | 1.0 | 4873 | 8383 |
|  | Spring mass | Autumn mass | 13.366 | 0.986 | 11.442 | 15.314 | 1.0 | 1853 | 3210 |
|  | Spring mass | Age. 1 | 2.562 | 2.818 | -3.065 | 8.005 | 1.0 | 2139 | 3735 |
|  | Spring mass | Age. 2 | 4.505 | 3.478 | -2.347 | 11.330 | 1.0 | 1930 | 3449 |
|  | Spring mass | Age. 3 | -3.323 | 2.868 | -9.035 | 2.242 | 1.0 | 2095 | 3790 |
|  | Spring mass | Population density | -0.950 | 0.561 | -2.064 | 0.155 | 1.0 | 8866 | 9970 |
|  | Spring mass | Average snow depth | -2.532 | 0.606 | -3.728 | -1.337 | 1.0 | 9347 | 10180 |
|  | Spring mass | Snow cover duration | -0.990 | 0.451 | -1.877 | -0.097 | 1.0 | 9804 | 11453 |
|  | Spring mass | Median snow density | 0.522 | 0.604 | -0.666 | 1.714 | 1.0 | 8820 | 11018 |
|  | Spring mass | Population density : Average snow depth | 0.755 | 0.669 | -0.561 | 2.080 | 1.0 | 10560 | 11409 |
|  | Spring mass | Population density : Snow cover duration | -0.220 | 0.528 | -1.242 | 0.816 | 1.0 | 9298 | 11036 |
|  | Spring mass | Population density : Median snow density | -1.214 | 0.606 | -2.401 | -0.030 | 1.0 | 10272 | 10601 |
|  | Next autumn mass | Spring mass | 17.664 | 0.466 | 16.747 | 18.581 | 1.0 | 5575 | 9863 |
|  | Next autumn mass | Age_t+1. 1 | -0.083 | 1.475 | -2.918 | 2.814 | 1.0 | 7819 | 11568 |
|  | Next autumn mass | Age_t+1. 2 | 0.468 | 1.749 | -2.984 | 3.901 | 1.0 | 5945 | 9706 |
|  | Next autumn mass | Age_t+1. 3 | 3.085 | 1.462 | 0.221 | 5.933 | 1.0 | 7730 | 11175 |
|  | Next autumn mass | Next population density | -0.559 | 0.568 | -1.678 | 0.572 | 1.0 | 4445 | 7023 |
